# Reconstitution of an intact clock that generates circadian DNA binding in vitro

**DOI:** 10.1101/2020.06.18.158576

**Authors:** Archana G. Chavan, Dustin C. Ernst, Mingxu Fang, Cigdem Sancar, Carrie L. Partch, Susan S. Golden, Andy LiWang

## Abstract

Circadian clocks control gene expression in the complex milieu of cells. Here, we reconstituted under defined conditions in vitro the cyanobacterial circadian clock system which includes an oscillator, signal-transduction pathways, transcription factor, and promoter DNA. The system oscillates autonomously with a near 24 h period, remains phase coherent for many days, and allows real-time observation of each component simultaneously without user intervention. This reassembled clock system provides new insights into how a circadian clock exerts control over gene expression and can serve in the area of synthetic biology as a new platform upon which to build even more complexity.

**One Sentence Summary:** An autonomously oscillating circadian clock-controlled gene regulatory circuit is studied in vitro using a real-time high-throughput assay.

---

Endogenous circadian clocks provide organisms with an internal representation of local time (*1*), and the *Synechococcus elongatus* clock generates *bona fide* circadian rhythms of genetic, physiological, and metabolic activities that fulfill all criteria that define circadian clocks (*2*). The core circadian clock genes of cyanobacteria, *kaiA, kaiB*, and *kaiC*, are essential for rhythmic gene expression (*3*), and their proteins generate an autonomous ∼24 h rhythm of KaiC phosphorylation in vivo (*4*) that can be recapitulated in vitro *(5)*. Ordered phosphorylation of KaiC at Thr 432 and Ser 431, as stimulated by KaiA (*6*) and suppressed by KaiB (*7*), helps set the pace of the core clock (*8, 9*). Genetic experiments have shown that this information is relayed to transcription machinery through signal-transduction pathways, but the standard in vitro KaiABC oscillator (*5*) did not provide a means to interrogate this process. Here, we report reconstitution of a clock-controlled gene regulation network in a cell-free system. This in vitro clock (IVC) includes the core KaiA, KaiB, and KaiC oscillator components, signal transduction enzymes (SasA and CikA), a transcription factor (RpaA), and a promoter-bearing DNA fragment (P*kaiBC*). We show that an IVC consisting of KaiA, KaiB, KaiC, RpaA, and SasA or CikA comprises the fundamental network needed for timekeeping, synchronization with the environment, and temporal relay to downstream events. Analysis of this network revealed steps in the clock mechanism that were not approachable previously.

Protein interactions in the IVC were monitored by real-time fluorescence anisotropy (*10*). KaiA, KaiB, SasA, CikA, and RpaA were fluorescently labeled using sortase-mediated ligation (*11, 12*). A fluorescently labeled synthetic dsDNA representing the promoter sequence of the *kaiBC* operon, P*kaiBC* (see Suppl. Mat. Table S2), was used as the RpaA target. Control experiments identified specific binding events that cause changes in the fluorescence anisotropies of each labeled component (Suppl. Mat. Fig. S1-S3). Using a multimode microplate reader, in vitro oscillations were monitored in parallel, with each reaction containing a single fluorescently labeled protein (50 nM) in addition to unlabeled proteins KaiA, KaiB, KaiC, SasA or CikA, and RpaA at physiological concentrations (*13*) (see Suppl. Mat. Tables S2 and S3 for details). Samples also contained either 100 nM fluorescently labeled or 100 nM unlabeled P*kaiBC* DNA. This protocol allowed measurement of the fluorescence anisotropies of each clock component and DNA in parallel reactions in a single plate and in real time over a duration of several days (Fig. 1A). The phosphorylation states of RpaA and KaiC were determined in parallel by gel electrophoresis (Suppl. Mat. Figs. S5 and S6), and the ATPase activity of KaiC was monitored in real time by ^1^H-NMR (Suppl. Mat. Fig. S7) (*14*). Data were fit to a cosine function to extract periods and relative phases (see Supplementary Material). Maximal KaiC phosphorylation was referenced to circadian time (CT) 14, which is when it peaks in vivo (*6*), and the relative phases of the other components are compiled in Fig. 1B. CT 0 and CT 12 represent subjective dawn and dusk under constant lab conditions.

**Fig. 1.**
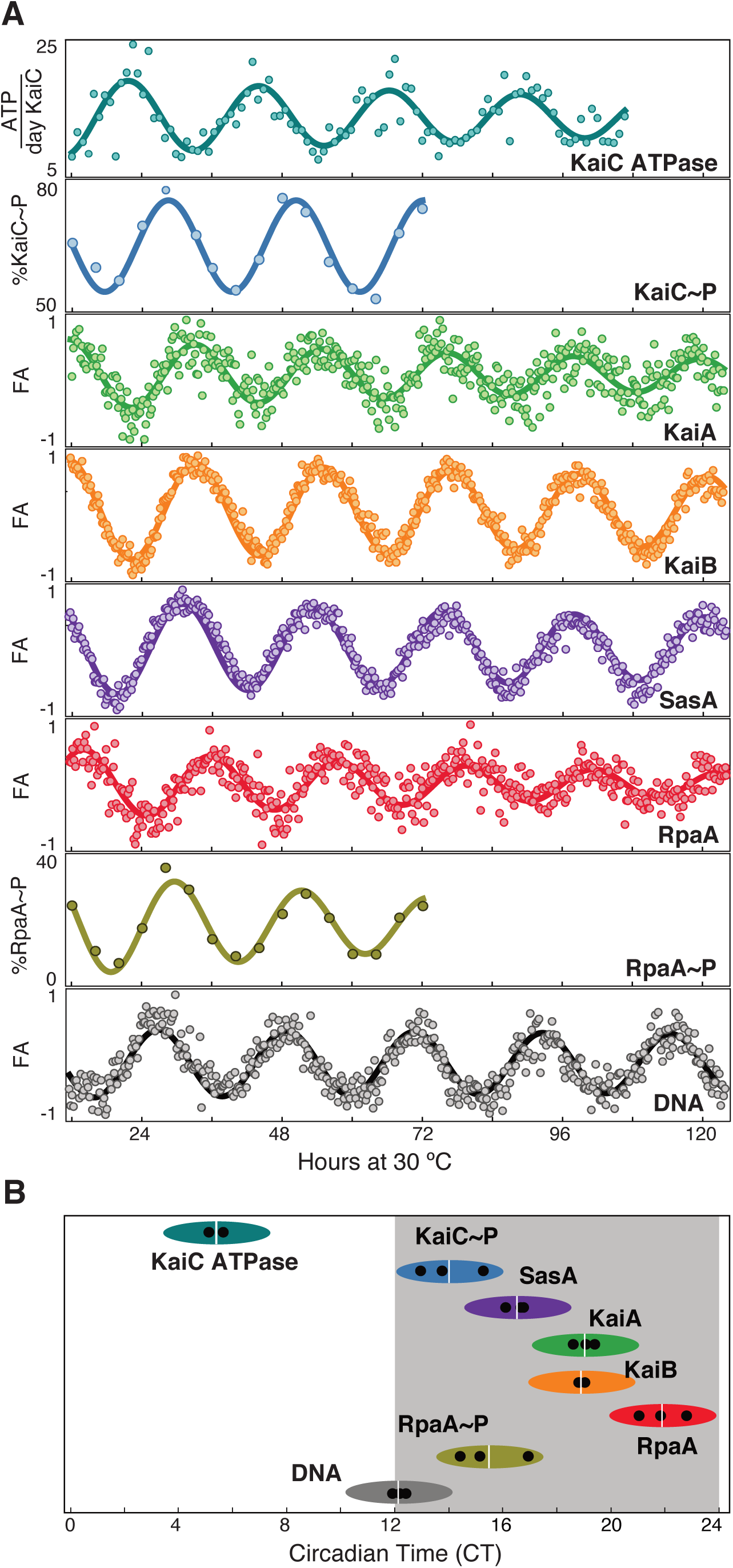
Reconstitution of an intact clock that controls DNA binding in vitro. (**A**) Clock reactions containing KaiA, KaiB, KaiC, SasA RpaA, DNA (P*kaiBC*), 1 mM ATP, and 5 mM MgCl_2_ were reconstituted in vitro. Fluorescence anisotropies (FA) for 50 nM probe KaiA (green), KaiB (orange), SasA (purple), RpaA (red) and 100 nM P*kaiBC* (gray), were measured in a microplate reader at 30 °C. For phosphorylation experiments 8 µL aliquots were collected every 4 hours and flash frozen in liquid N_2_, followed by gel electrophoresis. KaiC∼P (blue) was analyzed on 7.5% SDS-PAGE and RpaA∼P (olive) on 12.5% Zn^2+^-Phos-tag™ gels. KaiC-ATPase (teal) activity was measured by real-time ^1^H-NMR in a KaiABC-only reaction to isolate the KaiC ATPase activity. After truncating the first 12 hours of data collection during which samples approached stable limit cycles, data were baseline-corrected (filled circles) and fit to a cosine function (solid lines) to calculate periods and phases (see Materials and Methods). Fluorescence data were also normalized before fitting. **(B)** Relative phases of each component determined from three separate experiments (determined from cosine fits of data in (A)) are shown by black dots with the average denoted by a white vertical line. Peak of KaiC∼P is referenced to CT = 14 (*14*). The width of ellipses represents ≥80% peak height.

The relative phases of the in vitro rhythms in an IVC that contained KaiA, KaiB, KaiC, SasA, RpaA, and DNA are consistent with those reported in vivo (*2*). For example, fluorescence anisotropy rhythms of KaiA, KaiB, and SasA peaked around CT 15-22, indicating formation of large protein complexes during subjective night due to interactions involving KaiC (Fig. 1, Suppl. Mat. Fig. S1). In vivo these proteins are also found in large complexes around ZT 16-22 (*15*), where ZT 0 and ZT 12 are dawn and dusk under a light-dark cycle. SasA and KaiB compete for overlapping binding sites on KaiC (*16*) but SasA rhythms peaked ∼3 h earlier than those of KaiB as seen in Fig. 1B. This phase difference provides a temporal window during which SasA-KaiC interactions activate SasA autophosphorylation (*17*) allowing phosphoryl transfer from SasA to RpaA (*18*). Thus, RpaA phosphorylation and DNA binding reached their peaks around subjective dusk in the IVC. This earlier binding by SasA also assists subsequent KaiB-KaiC binding, extending the range of KaiB concentrations under which the oscillator is rhythmic as detailed in an accompanying report (Heisler and Swan et. al.). The fluorescence anisotropy of RpaA increases when it dephosphorylates or interacts with SasA in the absence of KaiC but decreases when it is phosphorylated by CikA or the SasA-KaiC-EE complex (Suppl. Mat. Figs. S2 and S3), which underlies the phases of RpaA seen in Figs 1 and 2. During subjective midday the ATPase activity of KaiC peaks at 13-18 ATP/KaiC monomer per day and during subjective night decreases to a nadir of 7-8 ATP/KaiC per day, consistent with earlier studies (*19*). Thus, the IVC recapitulates to a reasonable extent the relative in vivo phases of the circadian clock of *S*. *elongatus* and enables aspects of the clock — input (*20, 21*), oscillator (*5*), output (*13, 22–24*) — to be queried under highly defined conditions with multiple readouts in real time.

**Fig. 2.**
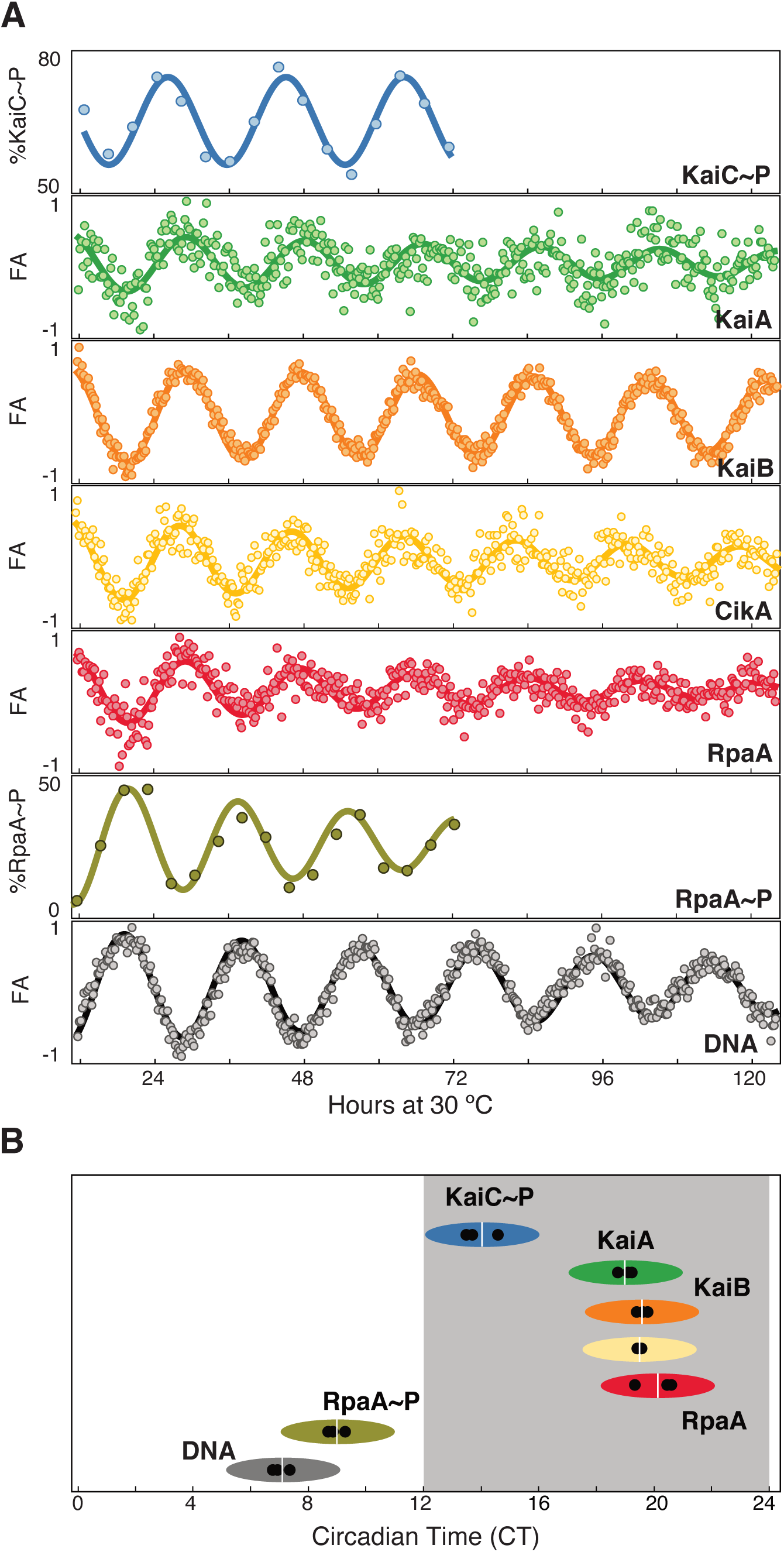
CikA transmits time signals downstream in the absence of SasA. **(A)** Fluorescence anisotropy rhythms of 50nM probe, KaiA (green), KaiB (orange), CikA (gold), RpaA (red), and 100 nM DNA (P*kaiBC*, gray) are shown along with KaiC∼P (blue) and RpaA∼P (olive) measured in clock reactions similar to those in Fig. 1A, but with CikA replacing SasA. **(B)** Relative phases of KaiC∼P, KaiA, and KaiB remain unchanged, while DNA peaks at midday (CT 7) instead of dusk (CT 12).

The standard model defines SasA as the cognate kinase of RpaA that activates gene expression at dusk (*25*), and the circadian input kinase CikA, important for phase resetting (*26*) as an RpaA phosphatase at night (*13, 24*). However, as shown in Fig. 2, when SasA was replaced by CikA in vitro, all clock components displayed robust circadian rhythms with KaiA, KaiB, and KaiC maintaining the same relative phases as in Fig 1B. The fluorescence anisotropy of CikA peaked with that of KaiB (Fig 2), consistent with the observation that the *pseudo*-receiver domain of CikA binds to the fold-switched state of KaiB that associates with KaiC (*27*). This interaction is known to stimulate the phosphatase activity of CikA (Suppl. Mat. Fig. S4) (*24, 28*); however, in the absence of this interaction with KaiB, CikA acts as a kinase toward RpaA (Suppl. Mat. Fig S4) (*24*). Hence, CikA is bifunctional and transduces clock signals antiphasic to those of SasA by switching between phosphatase and kinase activities, which may play a role in how it resets the clock in cyanobacteria under the application of dark pulses (*26*). Indeed, Δ*sasA* strains still generate circadian rhythms of bioluminescence in low light (*25*), possibly due to CikA (Suppl. Mat. Fig. S9), but Δ*sasA*Δ*cikA* double knockout strains are arrhythmic (*29*).

The IVC can provide insights into biological questions previously unapproachable by genetic studies, such as how clock control over RpaA regulates promoter activity. An *S*. *elongatus* strain expressing a single nucleotide polymorphism in an *rpaA* allele (originally reported as a *crm1* mutant (*30*)) resulted in an R121Q amino acyl substitution and did not produce rhythmic gene expression despite having a wild-type RpaA phosphorylation pattern (Suppl. Mat. Fig. S10). This arrhythmic allele *rpaA*-*R121Q* was reconstructed in a wild-type reporter strain (Fig. 3A) by markerless CRISPR-Cas12a editing (*31*). The particular transcription-activation step disrupted by the R121Q substitution in RpaA could not be predicted *a priori* (*32*). Using the IVC, we found that the RpaA R121Q mutant displays circadian rhythms of fluorescence anisotropy (Suppl. Mat. Fig. S11) and phosphorylation (Fig. 3B and Suppl. Mat. Fig. S12) like WT RpaA. However, the RpaA R121Q mutant bound the P*kaiBC* promoter poorly as indicated by the low amplitude of DNA fluorescence anisotropy rhythms. Thus, while the R121Q amino acyl substitution in RpaA does not hinder its interactions with SasA or CikA, the signal is not transduced normally to the DNA-binding domain, possibly by blocking phosphorylation-dependent oligomerization (*33*).

**Fig. 3.**
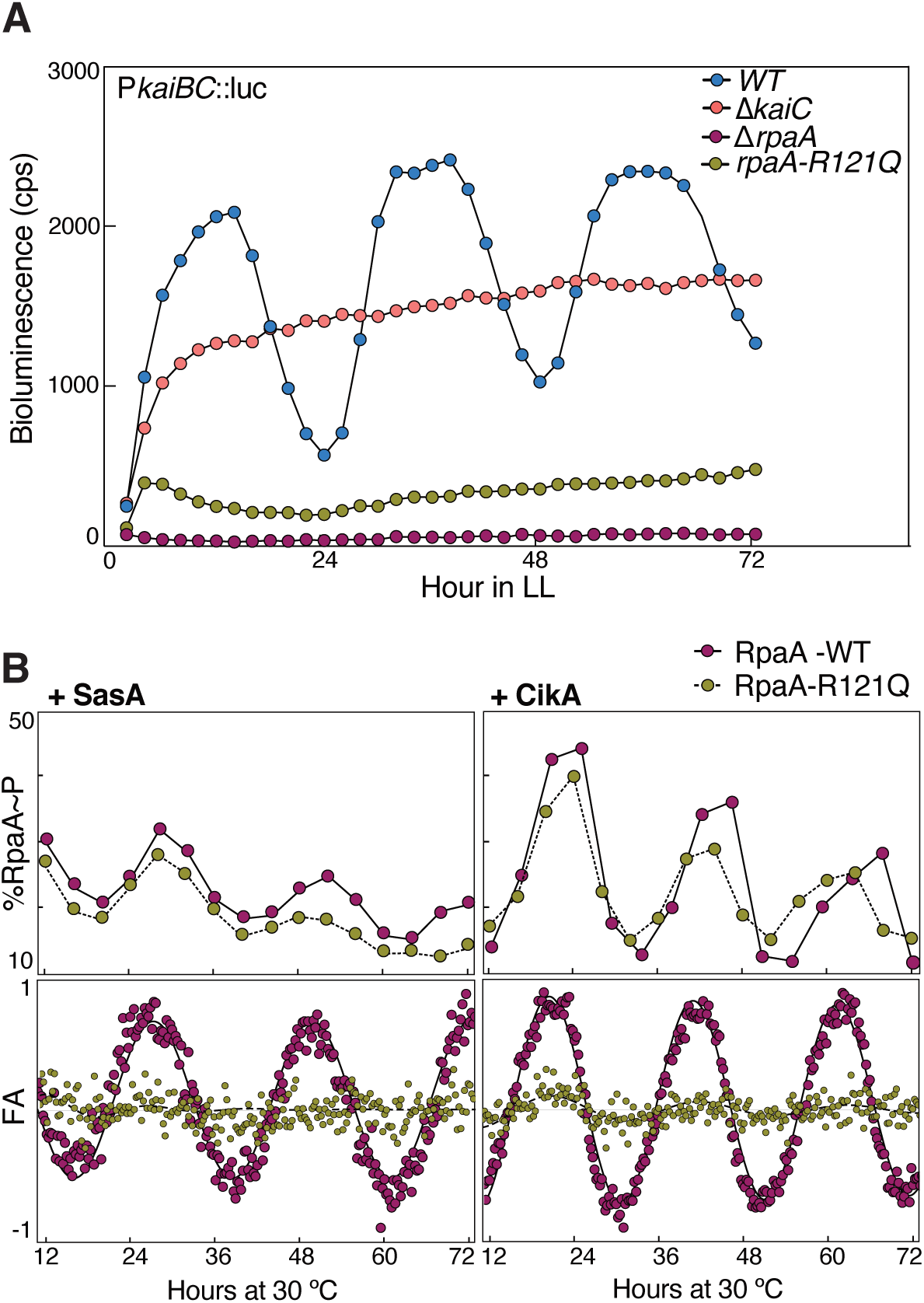
IVC explains why RpaA-R121Q mutant produces arrhythmic gene expression in cyanobacteria. **(A)** Rhythms of bioluminescence generated by P*kaiBC*::*luc* expression were monitored for entrained wild-type (blue), Δ*kaiC* (pink), Δ*rpaA* (maroon) and *rpaA*-R121Q (olive) cyanobacterial strains. Each plot represents an average of six biological replicates. **(B)** Top panels: Phosphorylation rhythms are for WT RpaA (maroon dots) and RpaA-R121Q mutant (olive dots) in in vitro reactions containing either SasA (left) or CikA (right). Bottom panels: Fluorescence anisotropy rhythms of DNA (P*kaiBC*) in the presence of WT RpaA (maroon dots) or RpaA-R121Q mutant (olive dots).

This extended in vitro clock is an autonomously oscillating system that remains phase coherent for several days without user intervention, allowing observation of rhythmic behaviors of several clock components simultaneously, including interactions with a promoter element on DNA. Complementing this assay with parallel measurements of phosphorylation and ATPase activities rounds out a moving picture of an intact cyanobacterial clock. The IVC now makes it possible to ask questions about how changing environments, such as temperature, metabolites, and protein levels are reflected in the core oscillator and propagated to regulation of transcription, providing deeper mechanistic insights into clock biology.

## Acknowledgments

We thank Joel Heisler, Jeffery Swan, Gary Chow, Yong-gang Chang, Michael Rust, Lu Hong, Shahar Sukenik, and Maria Zoghbi for helpful discussions. We also thanks to David Rice for maintaining the NMR Facility, and Steve Grimaldi for maintaining the NMR cryoplatform.

## Funding

This work was supported by US National Institutes of Health grants R01GM107521 (to A.L.), R01GM121507 (to C.L.P.) and R35GM118290 (to S.S.G.).

## Author contributions

A.L., A.G.C., D.C.E., M.F., C.S., C.L.P. and S.S.G. designed the experiments, and contributed reagents. A.G.C., D.E., M. F. and C. S. performed the experiments. Data were analyzed by A.G.C., A.L., D.E., M.F., C. S., C.L.P., and S.S.G. A.G.C., A.L., C.L.P., and S.S.G. wrote the paper. All authors approve of the conclusions and final version of the paper.

## Competing interests

Authors declare no competing interests.

## Data and materials availability

All data are available in the main text or the supplementary materials.

## Notes

### Competing Interest Statement

The authors have declared no competing interest.

