## Supplementary Materials for "Reconstitution of an intact clock that generates circadian DNA binding in vitro"

**This PDF file includes:**

Materials and Methods
Supplementary Text
Figs. S1 to S12
Tables S1 to S4

### Supplementary Information

### Materials and Methods

#### Cloning, expression and purification of proteins:

Clock genes from *Synechococcus elongatus* were cloned in pET-28b vector to produce 6x-His-SUMO fusion proteins in *Escherichia coli* BL21(DE3) (Agilent). A single colony from freshly transformed cells was used for inoculation of a starter culture grown in LB medium supplemented with 50 µg/mL kanamycin sulphate at 37 ºC, 220 rpm. After 6.5 h a 5 mL starter culture was transferred to 1 L M9 medium supplemented with 0.2% D-glucose, 2 mM $MgSO_4$, 0.1 mM $CaCl_2$, and 50 µg/mL kanamycin sulfate. Cells were grown to $OD_{600} \approx 0.6$ at 37 ºC before inducing expression by addition of 0.2 mM isopropyl β- d-1-thiogalactopyranoside (IPTG) and incubating cells at 30 ºC for 12 h except for CikA expression, for which cells were incubated at 20 ºC for 22 h.

Cells were harvested and lysed using an Avestin C3 Emulsiflex homogenizer (Avestin Inc, Canada). Cell lysate was clarified by high-speed centrifugation before affinity purification on Ni-NTA resin (QIAGEN) in polypropylene gravity columns using the buffers specified in Table 1.

All purification steps were carried out at 4 ºC. For cleavage of the 6x-His-SUMO fusion protein, 6x-His-ULP1 protease was added to final concentration of 3 µM and incubated at 4 ºC for 15 h. Reductant, tris(2-carboxyethyl) phosphine (TCEP) was added to SasA and CikA samples during ULP1 cleavage (concentrations given in Table S1 below). 6x-His-SUMO was removed by loading the cleaved protein on a Ni-NTA column for a second time. The flow-through was concentrated using 10 kDa molecular weight cut off (MWCO) membrane filters in an Amicon™ (Millipore Sigma) stir-celled concentrator at 4 ºC before purifying by gel-filtration chromatography.

**Table S1. Protein expression and purification summary**

| Protein construct | Expression temperature and duration | Ni-NTA Loading buffer | Ni-NTA Wash buffer | Ni-NTA Elution buffer | Gel Filtration Column and Elution Buffer |
|---|---|---|---|---|---|
| G-KaiA | 30 ºC, 12 h | 50 mM $NaH_2PO_4$, 500 mM NaCl, pH 8 | 50 mM $NaH_2PO_4$, 500 mM NaCl, 20 mM imidazole pH 8 | 50 mM $NaH_2PO_4$, 500 mM NaCl, 250 mM imidazole pH 8 | Superdex-200-1660-pg<br>20 mM Tris, 150 mM NaCl, 5 mM $MgCl_2$, 1 mM ATP, pH 8 |
| G-KaiB-FLAG | 30 ºC, 12 h | 50 mM $NaH_2PO_4$, 500 mM NaCl, pH 8 | 50 mM $NaH_2PO_4$, 500 mM NaCl, 20 mM imidazole pH 8 | 50 mM $NaH_2PO_4$, 500 mM NaCl, 250 mM imidazole pH 8 | Superdex-75-1660-pg<br>20 mM Tris, 150 mM NaCl, 5 mM $MgCl_2$, 1 mM ATP, pH 8 |
| FLAG-KaiC | 30 ºC, 12 h | 50 mM $NaH_2PO_4$, 500 mM NaCl, pH 8 | 50 mM $NaH_2PO_4$, 500 mM NaCl, 80 mM imidazole pH 8 | 50 mM $NaH_2PO_4$, 500 mM NaCl, 250 mM imidazole pH 8 | Superdex-200-1660-pg<br>20 mM Tris, 150 mM NaCl, 5 mM $MgCl_2$, 1 mM ATP, pH 8 |
| FLAG-KaiC-AA (S431A, T432A) | 30 ºC, 12 h | 50 mM $NaH_2PO_4$, 500 mM NaCl, pH 8 | 50 mM $NaH_2PO_4$, 500 mM NaCl, 80 mM imidazole pH 8 | 50 mM $NaH_2PO_4$, 500 mM NaCl, 250 mM imidazole pH 8 | Superdex-200-1660-pg<br>20 mM Tris, 150 mM NaCl, 5 mM $MgCl_2$, 1 mM ATP, pH 8 |
| FLAG-KaiC-AE (S431A, T432E) | 30 ºC, 12 h | 50 mM $NaH_2PO_4$, 500 mM NaCl, pH 8 | 50 mM $NaH_2PO_4$, 500 mM NaCl, 80 mM imidazole pH 8 | 50 mM $NaH_2PO_4$, 500 mM NaCl, 250 mM imidazole pH 8 | Superdex-200-1660-pg<br>20 mM Tris, 150 mM NaCl, 5 mM $MgCl_2$, 1 mM ATP, pH 8 |
| FLAG-KaiC-EE (S431E, T432E) | 30 ºC, 12 h | 50 mM $NaH_2PO_4$, 500 mM NaCl, pH 8 | 50 mM $NaH_2PO_4$, 500 mM NaCl, 80 mM imidazole pH 8 | 50 mM $NaH_2PO_4$, 500 mM NaCl, 250 mM imidazole pH 8 | Superdex-200-1660-pg<br>20 mM Tris, 150 mM NaCl, 5 mM $MgCl_2$, 1 mM ATP, pH 8 |

| | | | | | |
|---|---|---|---|---|---|
| FLAG-KaiC-EA (S431E, T432A) | 30 ºC, 12 h | 50 mM $NaH_2PO_4$, 500 mM NaCl, pH 8 | 50 mM $NaH_2PO_4$, 500 mM NaCl, 80 mM imidazole pH 8 | 50 mM $NaH_2PO_4$, 500 mM NaCl, 250 mM imidazole pH 8 | Superdex-200-1660-pg<br>20 mM Tris, 150 mM NaCl, 5 mM $MgCl_2$,1 mM ATP, pH 8 |
| G-SasA | 30 ºC, 12 h | 50 mM $NaH_2PO_4$, 500 mM NaCl, pH 8 | 50 mM $NaH_2PO_4$, 500 mM NaCl, 20 mM imidazole pH 8 | 50 mM $NaH_2PO_4$, 500 mM NaCl, 250 mM imidazole pH 8 + 10 mM TCEP | Superdex-200-1660-pg<br>20 mM Tris, 150 mM NaCl, 5 mM $MgCl_2$, 1 mM ATP, pH 8 |
| G-SasA-H161A | 30 ºC, 12 h | 50 mM $NaH_2PO_4$, 500 mM NaCl, pH 8 | 50 mM $NaH_2PO_4$, 500 mM NaCl, 20 mM imidazole pH 8 | 50 mM $NaH_2PO_4$, 500 mM NaCl, 250 mM imidazole pH 8 + 10 mM TCEP | Superdex-200-1660-pg<br>20 mM Tris, 150 mM NaCl, 5 mM $MgCl_2$, 1 mM ATP, pH 8 |
| FLAG-CikA-LPETGG | 20 ºC, 22 h | 50 mM $NaH_2PO_4$, 500 mM NaCl, pH 8 | 50 mM $NaH_2PO_4$, 500 mM NaCl, 20 mM imidazole pH 8 | 50 mM $NaH_2PO_4$, 500 mM NaCl, 250 mM imidazole pH 8 + 20 mM TCEP | Superdex-200-1660-pg<br>20 mM Tris, 150 mM NaCl, 5 mM $MgCl_2$, pH 8 |
| GGG-RpaA | 30 ºC, 12 h | 50 mM $NaH_2PO_4$, 500 mM NaCl, pH 8 | 50 mM $NaH_2PO_4$, 500 mM NaCl, 20 mM imidazole pH 8 | 50 mM $NaH_2PO_4$, 500 mM NaCl, 250 mM imidazole pH 8 | Superdex-75-1660-pg<br>20 mM Tris, 150 mM NaCl, 5 mM $MgCl_2$, 1 mM ATP, pH 8 |
| GGG-RpaA-R121Q | 30 ºC, 12 h | 50 mM $NaH_2PO_4$, 500 mM NaCl, pH 8 | 50 mM $NaH_2PO_4$, 500 mM NaCl, 20 mM imidazole pH 8 | 50 mM $NaH_2PO_4$, 500 mM NaCl, 250 mM imidazole pH 8 | Superdex-75-1660-pg<br>20 mM Tris, 150 mM NaCl, 5 mM $MgCl_2$, 1 mM ATP, pH 8 |

#### Fluorescent labeling of proteins by sortase-mediated ligation:

N-terminal sortase-mediated ligations were performed for protein constructs with an N-terminal glycyl residue. The fluorophore-containing peptides 5-carboxyfluorescein (5-FAM)-LPETGG and C-(Cy3)-LPETGG were purchased from GenScript (New Jersey). For C-terminal ligations, target proteins were fused to sortase-A recognition peptide LPETGG at the C-terminus. The peptide GGGYCN was expressed and purified in-house as a 6x-His-SUMO-fusion protein. The cysteinyl residue on the purified fusion protein was labeled with the fluorophore 6-iodoacetamidofluorescein (6-IAF) (Invitrogen) as described previously (*1*). The labeled peptide was cut from the SUMO-fusion protein using ULP1 and passed through 10 kDa MWCO membrane in stirred cell concentrator. The flow through containing labeled peptide was further purified by C4-reverse-phase column chromatography. The peptide eluted with a linear acetonitrile gradient was then lyophilized. Dried peptide was dissolved in sortase buffer before use. Concentration was determined by UV absorbance at 493 nm for 6-IAF fluorophore.

Each target protein was buffer exchanged in sortase buffer (20 mM Tris, 150 mM NaCl, pH 7.5) before ligation. For ligation, 50 μM target protein was incubated with 250 μM fluorophore peptide, 5 μM sortase-A, and 10 mM $CaCl_2$ in the dark at 4 ºC for 12-16 h. Sortase-A was separated from reaction mixtures by Ni-NTA chromatography using prepacked 5 mL HiTrap-Ni-NTA columns (GE Healthcare) on an AKTA FPLC using a step gradient of imidazole. Labeled protein fractions were concentrated and purified further by gel-filtration chromatography using a Superose-6-Increase-10/300 GL analytical grade column (GE Healthcare). Table S2 lists the fluorophore peptides used for labeling each protein construct.

**Table S2. Fluorophore labels and target proteins ligated using sortase-A**

| Target Protein | Label Position | Peptide with Fluorophore | Fluorophore Excitation Max (nm) | Fluorophore Emission Max (nm) |
|---|---|---|---|---|
| G-KaiA | N-terminal | C(Cy3)-LPETGG | 554 | 568 |
| G-KaiB-FLAG | N-terminal | 5-FAM-LPETGG | 480 | 520 |
| G-SasA | N-terminal | 5-FAM-LPETGG | 480 | 520 |
| G-SasA-H161A | N-terminal | 5-FAM-LPETGG | 480 | 520 |
| GGG-RpaA | N-terminal | 5-FAM-LPETGG | 480 | 520 |
| GGG-RpaA-R121Q | N-terminal | 5-FAM-LPETGG | 480 | 520 |
| FLAG-CikA-LPETGG | C-terminal | GGGYC(6-IAF)-N | 493 | 520 |
| **Fluorescently labeled synthetic DNA (Integrated DNA Technologies, Inc., Iowa, USA)**<br>(/4iCy3/ – internal labeling of DNA backbone with cyanoethyl phosphoramidite chemistry) | | | | |
| **P*kaiBC* 31-bp promoter sequence**<br>5'-CCG/4iCy3/AGC TTA AGA CCT CCT TTA CCT TTT CAG G-3'<br>3'-GGC TCG AAT TCT GGA GGA AAT GGA AAA GTC C-5' | | | 550 | 564 |

#### Reconstitution of clock reaction for fluorescence anisotropy measurement:

For monitoring multiple probes simultaneously, a master mix of oscillator reactions was prepared in 20 mM Tris, 150 mM NaCl, 5 mM $MgCl_2$, 1 mM ATP, and pH 8.0 buffer by mixing unlabeled proteins at the final concentrations shown in Table S3. Aliquots (95 µL) of reaction master mix were pipetted into black, non-binding 384-well microplates (Greiner BioOne). A 5 µL- aliquot of fluorophore-labeled protein (50 nM final concentration) or DNA (100 nM final concentration) was added to each well (and mixed by pipetting up and down. The plate was sealed using clear, adhesive film (MicroAmp$^{TM}$). The plate was transferred to a CLARIOstar multimode microplate reader (BMG Labtech), pre-equilibrated at 30 ºC. The plate was incubated for 30 minutes prior to starting measurements to allow equilibration of all samples. Detector gain for parallel and perpendicular channels was adjusted on a 50 nM fluorescent probe mixed in buffer, where the target polarization was set to 10% of the theoretical polarization maximum of fluorescein (350 mili Polarization units). Fluorescence polarization was measured in kinetic mode, every 15 minutes for 800 cycles. Fluorescence polarization (FP) was converted to fluorescence anisotropy (FA) in MARS software before analysis. All the anisotropy values are reported in mili anisotropy (mA) units.

**Table S3. In vitro clock reaction composition.**

| Component | Concentration |
|---|---|
| KaiA | 1.20 µM |
| KaiB-FLAG | 3.50 µM |
| FLAG-KaiC | 3.50 µM |
| SasA | 0.65 µM |
| FLAG-CikA | 0.65 µM |
| RpaA | 2.50 µM |
| P*kaiBC,*31-bp promoter DNA | 100 nM |

#### Phosphorylation assay for KaiC and RpaA

For phosphorylation measurements 250 µL samples from identical clock reactions were incubated in parallel with the fluorescence anisotropy experiments. For detection of RpaA phospho-states, fluorophore-labeled RpaA (50% of total concentration) was used. Samples were incubated at 30 ºC in a benchtop water bath. Aliquots (8 µL) were removed every 4 h by manual pipetting over a period of 72 h. Each aliquot was immediately flash frozen by dipping the tubes in liquid nitrogen and stored at -80 ºC until all the aliquots were collected. For phosphorylation analysis, time points were first run on $Zn^{2+}$-Phos-tag$^{TM}$ gels as described below and then the same set of aliquots was run on 7.5% SDS-PAGE for analysis of KaiC~P states.

***Sample preparation and analysis on Phos-tag$^{TM}$ gel:*** Frozen protein aliquots were thawed on ice for 15 minutes and mixed with 8 µL of chilled 2x-SDS-PAGE loading buffer. Samples were spun at 1500 rpm for 30 seconds and transferred quickly to ice. In each lane 4 µL sample was loaded on precast 50 µM-$Zn^{2+}$-Phos-tag$^{TM}$ -12.5% SDS-PAGE gels (Wako Chemicals, Japan). Phosphorylated RpaA was separated by electrophoresis at constant voltage (140 V) for 2 h in freshly prepared 1x running buffer (100 mM Tris, 100 mM MOPS, 0.1% SDS and 5 mM sodium bisulfite, pH 7.8). The electrophoresis apparatus was kept in an ice bath during electrophoresis and running buffer was chilled to 4 ºC before use. Fluorescent bands of RpaA were visualized under UV transillumination (E-Gel Imager, Thermo fisher) immediately after electrophoresis was completed and before staining the gels with CBB.

***SDS-PAGE for KaiC phosphorylation:*** Aliquots prepared for Phos-tag™ gels (above) were saved at -20 ºC until analysis. Samples were boiled at 95 ºC for 5 min and spun down at 15000 rpm for 30 seconds before loading on 7.5% SDS-PAGE gels. Samples (4 μL) were loaded in each well and two-step electrophoresis was performed at 60 V for 30 min followed by at 140 V for 1 h 40 min. The electrophoresis apparatus was kept in ice bath and pre-chilled running buffer was used. Gels were stained with InstantBlue® protein gel stain (Expedeon Inc., Novus Biologicals, Centennial Co.) for at least 1 h followed by de-staining in D.I. water for 30 minutes. Gels were imaged by E-gel Imager (Invitrogen, Thermo Fisher Scientific). Densitometry of gel images was performed using NIH ImageJ software (*2*).

#### ATPase activity measured by 1H-NMR

A solution of 1.2 μM KaiA, 3.5 μM KaiB, 3.5 μM KaiC, 95% $H_2O$, 5% $D_2O$, and 10 μM DSS was prepared in a reaction buffer containing 20 mM Tris, 100 mM NaCl, 1 mM ATP, 5 mM $MgCl_2$, pH 8. One-dimensional proton NMR spectra were measured at 30 ºC, every hour for 5 days. The ATP and ADP peak were fit using an interpolation function in Wolfram Mathematica. Peak intensities were plotted as function of time and fit to a straight line from which overall ATPase activity was determined. Oscillations of ATP and ADP resonances about the line provided the determination of ATPase rhythms (see Figure S6 for details).

#### Data fitting for phase and period analysis

First 12 h of raw data were removed before analysis because during this time samples are approaching stable limit cycles. Anisotropy and phosphorylation data were baseline corrected using quadratic function and normalized to +/-1 before fitting to a single cosine function:

$$Y = a\cos\left[2\pi\left(\frac{t}{p}\right) - \phi\right]e^{-kt}$$

where, $p$ = *period*, $\phi = phase\ angle$, $k$ = *decay constant.*

#### Strains and culture conditions

*Synechococcus elongatus* PCC 7942 and its derivative strains (Table S4) were maintained on BG-11 medium containing antibiotics as needed for selection (*3*). Growth on plates or in liquid medium was carried out at 30 ˚C under 150 µmol m$^{-2}$ s$^{-1}$ light. *E. coli* DH5α used for cloning was grown on LB with the appropriate antibiotics at 37 ˚C.

**Table S4.** ***Synechococcus elongatus*** **PCC7942 strains used for in vivo experiments.**

| **Strain** | **Genotype** | **Source** |
|---|---|---|
| AMC541 | WT. *S. elongatus* PCC 7942 / NSII-P*kaiBC::luc* | Lab collection |
| AMC704 | Δ*kaiC* / NSII-P*kaiBC::luc* | Lab collection |
| AMC1722 | Tn*5* 8S15-E11 in AMC541 | Boyd et al., 2013 (*4*) |
| Δ*rpaA* | *rpaA*::Gm / NSII-P*kaiBC::luc* | This study |
| *rpaA-R121Q* | *rpaA*(G362->A) / NSII-P*kaiBC::luc* | This study |
| **Plasmid** | **Description** | |
| pSL2680 | RSF1010-backbone, Cas12a and CRISPR assay from *Francisella novicida*, Km$^R$ | Ungerer and Pakrasi, 2016 |
| AM4523 | Deletion construct containing *rpaA*::Gm flanked by *S. elongatus* gDNA | Lab collection |
| pDE32 | pSL2680 with gRNA targeting *rpaA* and HDR encoding *rpaA* (G362->A) | This study |
| **Primer** | **Sequence** | |
| rpaA_gRNA_F | AGATGCTGAACAGCTAAAGCCTGA | This study |

| | | |
|---|---|---|
| rpaA_gRNA_R | AGACTCAGGCTTTAGCTGTTCAGC | This study |
| rpaA_HDR-UP_F | CATTTTTTTGTCTAGCTTTAATGCGGTAGTTGGTACCG AGTCCTGAGCTGCTACTGCC | This study |
| rpaA_HDR-UP_R | GGTCAGGCTTTAGCTGTGCAGCTCGTTCCCGACCTGA T | This study |
| rpaA_HDR-DWN_F | ATCAGGTCGGGAACGAGCTGCACAGCTAAAGCCTGA CC | This study |
| rpaA_HDR-DWN_R | GCCCGGATTACAGATCCTCTAGAGTCGACGGTACCGC CATGTCAAACTCAATCAAGCG | This study |
| rpaA_cPCR_F | GAACAGGCTGAAACGATGGC | This study |
| rpaA_cPCR_R | GGATTGAGTAATTTGATGGTTGACTGC | This study |

***Construction of rpaA-R121Q strain***: Introduction of point mutations into the *S. elongatus* chromosome was accomplished using a previously described CRISPR-editing approach (*5*). The pSL2680 ($Km^R$) plasmid used for CRISPR-Cas12a (formerly Cpf1) editing was purchased from Addgene (Plasmid #85581). Primers rpaA_gRNA_F and rpaA_gRNA_R were annealed together and ligated into AarI-cut pSL2680 to serve as the *rpaA-R121Q* gRNA template. The resulting construct was purified and digested with KpnI to facilitate insertion of the *rpaA-R121Q* homology directed repair (HDR) template. The HDR template was generated by amplifying overlapping upstream and downstream fragments using primers rpaA_HDR-UP_F and rpaA_HDR-UP_R (AMC1722 genomic DNA as template) and rpaA_HDR-DWN_F and rpaA_HDR-DWN_R (AMC541 genomic DNA as template), respectively. The upstream and downstream HDR fragments were assembled into KpnI-cut pSL2680+gRNA using the GeneArt Seamless Assembly Kit (Thermo Fisher Scientific), forming pDE32.

Plasmid pDE32 was electroporated into *E. coli* DH10B containing a helper plasmid pRL623 (chloramphenicol resistance, $Cm^R$) and conjugal plasmid pRL443 (ampicillin resistance, $Ap^R$) (*6*). The resulting strain was grown overnight in LB medium containing antibiotics (Ap, kanamycin Km, and Cm), washed 3x with fresh LB, and mixed in a 1:2 ratio with an *S. elongatus* reporter-strain aliquot. The cell mixture was plated onto BG-11 agar with added LB (5% vol/vol), incubated under 100 μmol $m^{-2}$ $s^{-1}$ light for 36 h, then underlaid with Km (10 μg/ml final concentration) to select for *S. elongatus* cells that contain pDE32. Colonies that emerged after 6-8 days were passaged three times on BG-11 agar containing Km to allow editing to occur. Successful editing of chromosomal *rpaA* was verified by sequencing. Plasmid pDE32 was cured from the edited strain by inoculating cells into non-selective BG-11 medium, growing the culture to $OD_{750}$ = 0.6, then dilution plating on non-selective BG-11 plates. Fifty colonies were picked and replica patched to selective (Km) and non-selective medium to identify and isolate clones that had lost pDE32.

***Bioluminescence Monitoring:*** Bioluminescence was monitored using a P*kaiBC*::*luc* firefly luciferase fusion reporter inserted into a neutral site of the *S. elongatus* chromosome as previously described (*3*). Strains to be monitored were grown in liquid culture to $OD_{750}$ = 0.4-0.7, diluted to $OD_{750}$ = 0.2, and added as 20 µl aliquots to 280 µl of BG-11 agar containing 3.5 mM firefly luciferin arrayed in 96-well plates. Plates were covered with a gas-permeable seal and cells were entrained under 12-h light-dark cycles (80 µmol $m^{-2}$ $s^{-1}$ light) to synchronize clock phases. After 48 h of entrainment, cells were released into continuous light (30 µmol $m^{-2}$ $s^{-1}$) and bioluminescence was monitored every 2 h using a Tecan Infinite Pro M200 Bioluminescence Plate Reader. Data were collected and plotted using GraphPad Prism 8, with each plot representing the average of six biological replicates.

***Immunoblotting:*** Flask-grown cells were collected (15 ml) at ZT 0 and ZT 12 from liquid cultures ($OD_{750}$ = 0.6-0.7) incubated under 12-h light-dark cycles (40 µmol $m^{-2}$ $s^{-1}$). Cells were pelleted, washed once with cold 10 mM sodium chloride solution and frozen at -80˚C. Cell pellets were thawed on ice and resuspended in tris-buffered saline (pH 7.4) containing 1 mM phenylmethylsulfonyl fluoride, then disrupted by bead beating at 4 ˚C (30 sec of beating, followed by 2 min on ice for 10 cycles). Following centrifugation (20,000 x *g* for 10 min), protein concentrations were determined by the Bradford assay and a total of 10 µg of protein was loaded per well.

Phos-tag$^{TM}$ reagent (25 µM) (Wako Chemicals, Japan) and manganese chloride (50 mM) were added to standard SDS-PAGE gels (12.5 %) to allow detection of phosphorylated RpaA. Electrophoresis was conducted on ice using pre-chilled running buffer to limit hydrolysis of the heat-labile phospho-aspartate. Current was maintained at 25 mA until the bromophenol blue dye reached the bottom edge of the gel. The gel was then incubated for 10 min in transfer buffer containing 10 mM EDTA, followed by a 10 min incubation in transfer buffer without EDTA prior to semi-dry transfer to a PVDF membrane using a Trans-Blot Turbo System (BioRad). Detection of RpaA was achieved using RpaA-antiserum (gift from E. O'Shea, HHMI–Janelia, Ashburn, VA) at a dilution of 1:2000 as described previously (*7*, *8*). Secondary antibody (goat anti-rabbit IgG; 401315, Calbiochem) was used at a dilution of 1:100,000 and chemiluminescent signal was produced using the SuperSignal West Femto Maximum Sensitivity Substrate (Thermo Fisher Scientific).

#### Supplementary Data and Figures

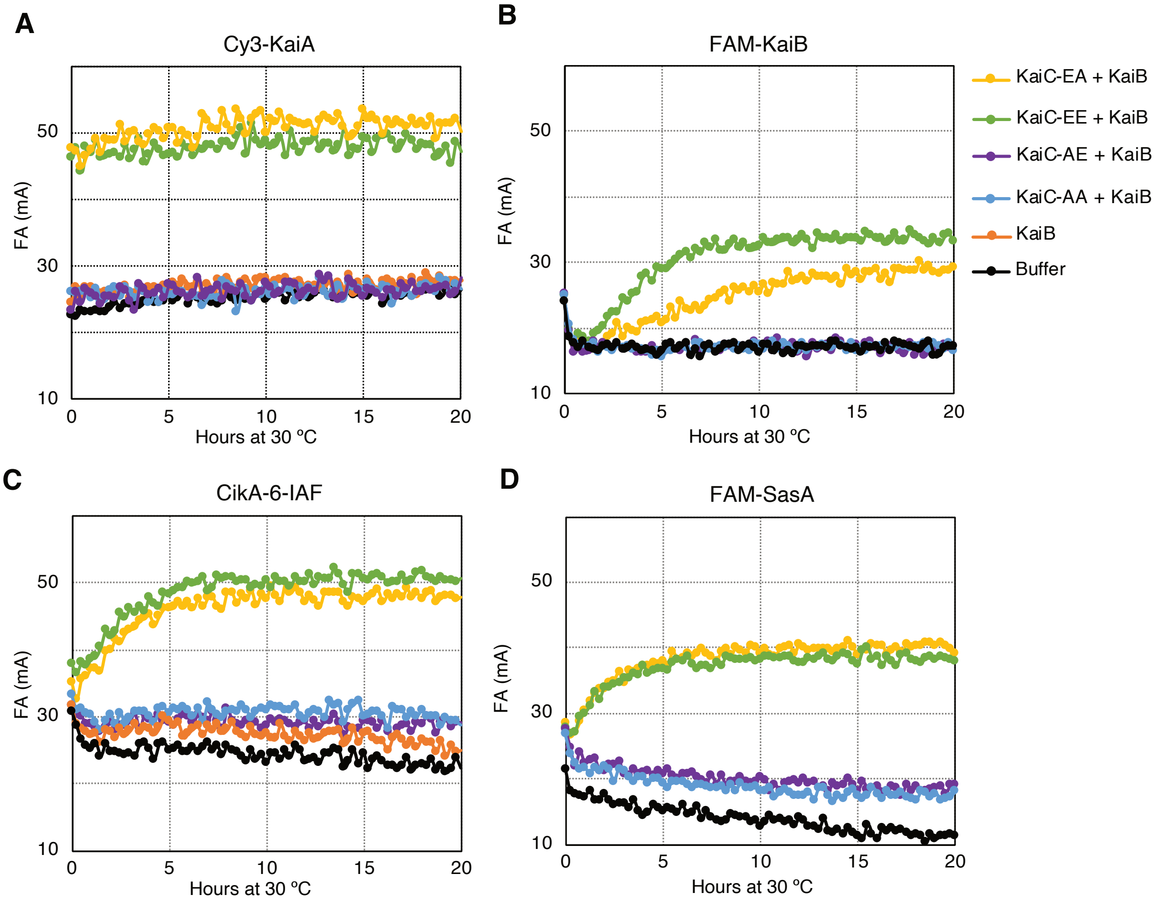

**Figure S1. Fluorescence anisotropy of KaiA, KaiB, CikA, and SasA in partial clock reactions.**

Control experiments of fluorescence anisotropies (FA) of labeled clock proteins at 50 nM were incubated with 3.5 µM KaiB (orange), or a mixture of each KaiC phosphomimetic (3.5 µM) with KaiB (3.5 µM). Anisotropies of each labeled protein mixed with buffer are shown in black. The four phospho-states of KaiC can be mimicked by mutating S431 and T432 to either Ala or Glu (*9*). These phosphostates are indicated as KaiC-AA (S431A, T432A, blue), KaiC-AE (S431A, S432E, violet), KaiC-EE (S431E, T432E, green) and KaiC-EA (S431E, T432A, yellow). KaiC-EE and KaiC-EA approximate the dusk pS,pT and nighttime pS,T states of KaiC. **(A)** Cy3-KaiA: G-KaiA was labeled at the N-terminus with C(-Cy3)-LPETGG. **(B)** FAM-KaiB: G-KaiB was labeled at the N-terminus with 5-FAM-LPETGG **(C)** CikA-6-IAF: CikA-LPETG was labeled at the C-terminus with GGGYC-(6-IAF)N, and **(D)** FAM-SasA: G-SasA was labeled at the N-terminus with 5-FAM-LPETGG.

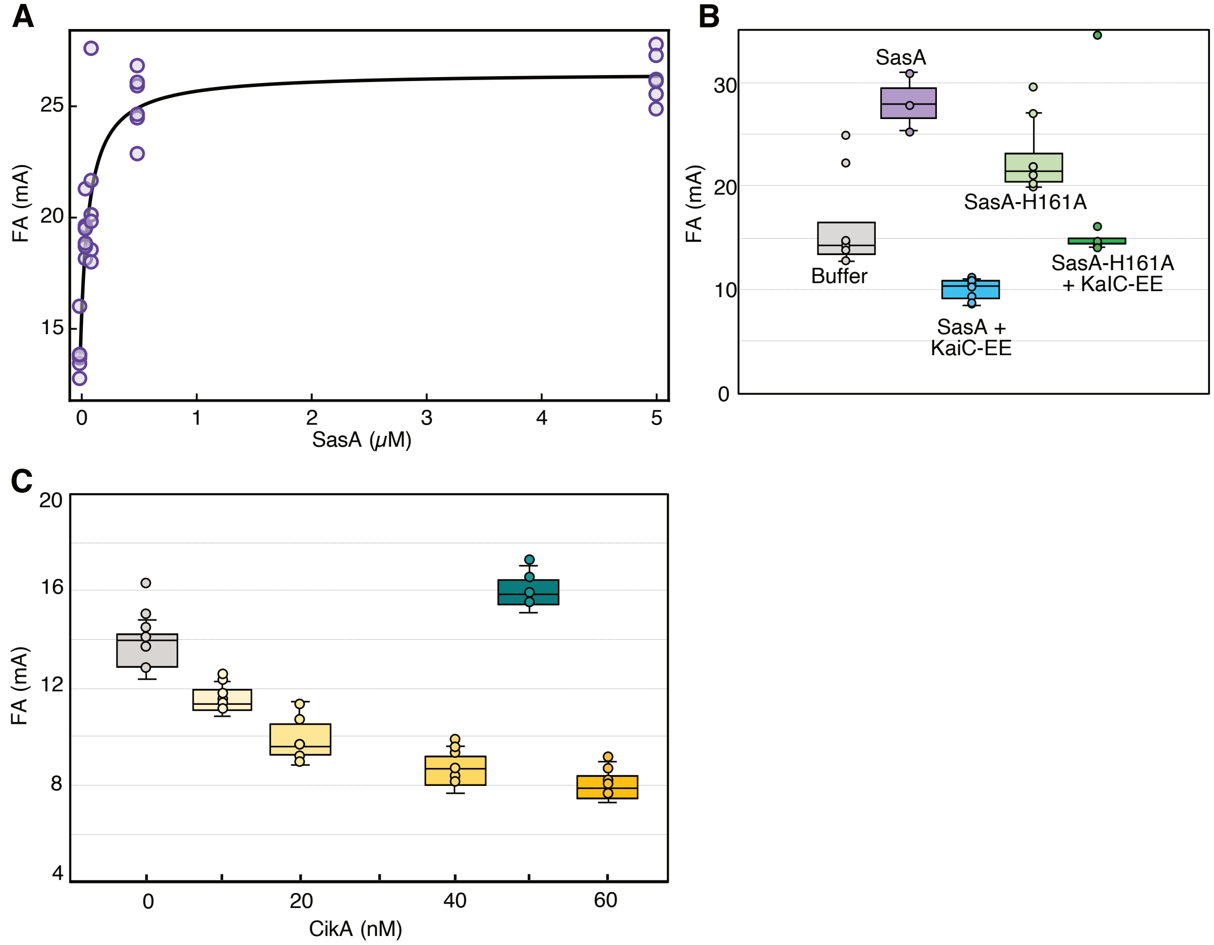

#### Figure S2. Fluorescence anisotropy of RpaA in the presence of SasA and CikA.

FAM-labeled RpaA (50 nM) was incubated with varying amounts of SasA **(A)** or CikA **(C)** and fluorescence anisotropy was measured at 30 ºC in kinetic mode, every 15 minutes for 12 h. Reported anistropies are averages of last 2 h of measuement which is when all the reactions reached steady state**. (A)** Fluorescence anisotropy of FAM-labeled RpaA (50 nM) from six replicates is plotted as a function of SasA concentration (purple circles) and data were fit to the following equation (*10*):

$$FA = FA_i + (FA_{max} - FA_i)\left(\frac{[SasA]}{[SasA] + K_D}\right)$$

where, FA = fluorescence anisotropy of RpaA,

$FA_i$ = initial fluorescence anisotropy of RpaA,

$FA_{max}$ = maximum fluorescence anisotropy of RpaA,

[SasA] = concentration of SasA (μM),

$K_D$ = apparent dissociation constant (μM).

Fitting the six data sets separately, the average apparent dissociation constant ($K_D$) is 0.07 ± 0.02 μM. **(B)** Box-whiskers plot for fluorescence anisotropies from six replicates of RpaA alone (gray) when incubated with 0.65 μM SasA (purple, 3 replicates), 0.65 μM SasA-H161A mutant (light green), 0.65 μM SasA + 3.5 μM KaiC-EE (blue), or 0.65 μM SasA-H161A + KaiC-EE (dark green). **(C)** Box-whiskers plot for RpaA fluorescence anisotropies measured from six replicates as a function of CikA concentration, and when incubated with 50 nM of CikA-H393A (teal, 4 replicates).

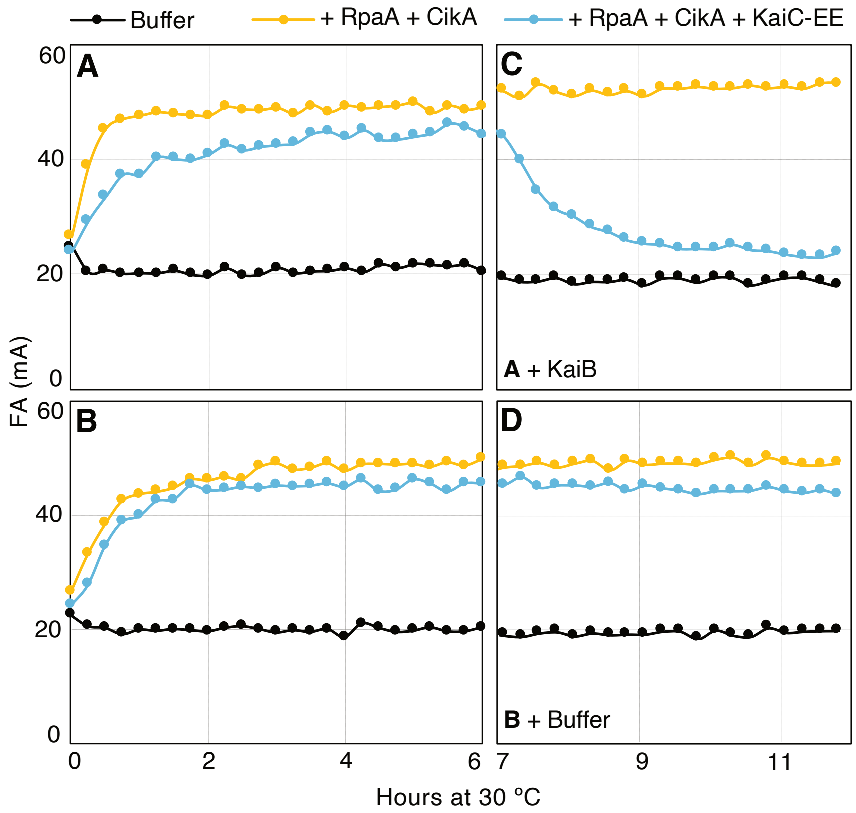

**Figure S3. Fluorescence anisotropies of DNA in the presence of clock components.**

Fluorescence anisotropies of Cy3-labeled P*kaiBC* DNA (100 nM) was measured in buffer containing 1 mM ATP and 5 mM $MgCl_2$, pH 8. (**A**) P*kaiBC* alone (black), P*kaiBC* + RpaA+ CikA (yellow), and P*kaiBC* + RpaA+ CikA + KaiC-EE (light blue). (**B**) Replicate of (A). (**C**) KaiB was added to samples in (A). (**D**) Buffer was added to samples in (B). After addition of KaiB or buffer at $t$ = 6 h, samples were incubated for 1 h before measuring anisotropies for another 5 h. Sample concentrations were 2.5 μM (RpaA), 0.65 μM (CikA), 3.5 μM (KaiC-EE), and 3.5 μM (KaiB).

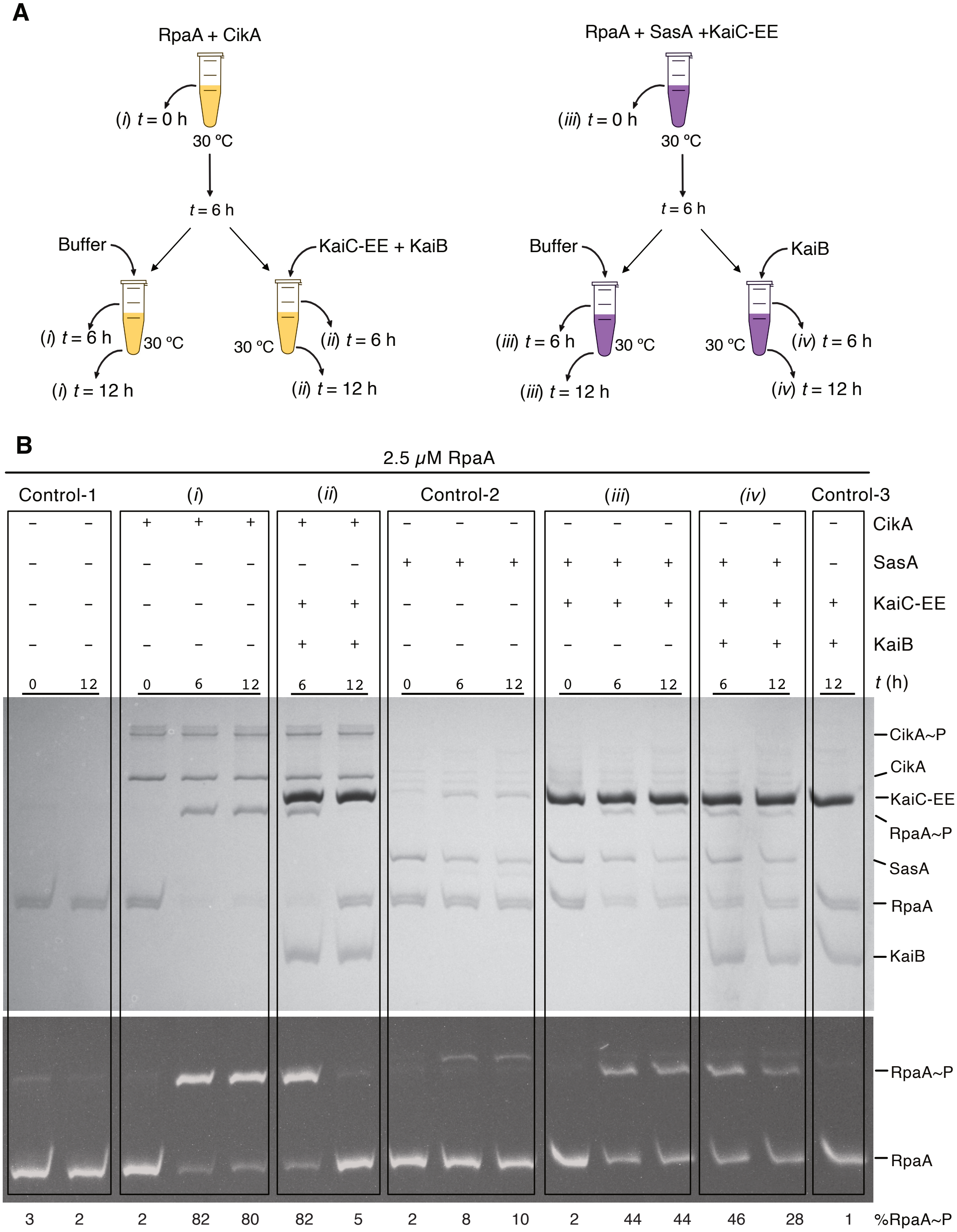
A
RpaA + CikA
(i) t = 0 h
30 °C
t = 6 h
Buffer
KaiC-EE + KaiB
(i) t = 6 h
30 °C
(i) t = 12 h
(ii) t = 6 h
(ii) t = 12 h
RpaA + SasA +KaiC-EE
(iii) t = 0 h
KaiB
(iii) t = 6 h
(iii) t = 12 h
(iv) t = 6 h
(iv) t = 12 h
B
2.5 μM RpaA
Control-1
(i)
(ii)
Control-2
(iii)
(iv)
Control-3
CikA
SasA
KaiC-EE
KaiB
t (h)
CikA~P
CikA
KaiC-EE
RpaA~P
SasA
RpaA
KaiB
%RpaA~P
3 2 2 82 80 82 5 2 8 10 2 44 44 46 28 1

#### Figure S4. RpaA phosphorylation assay in partial clock reactions.

**(A)** Schematics for RpaA phosphorylation assay in the presence of CikA (left panel) and SasA + KaiC-EE (right panel) ± KaiB. FAM-RpaA (2.5 µM) was incubated with either CikA (0.65 µM) or a mixture of SasA (0.65 µM) and KaiC-EE (3.5 µM) at 30 ºC in a buffer containing 1 mM ATP and 5 mM $MgCl_2$, pH 8. In the left panel an aliquot was removed immediately after RpaA and CikA were mixed (*t* = 0 h). The reaction was incubated at 30 ºC for 6 h then divided into two equal portions, one portion received an equal volume of buffer and other received a pre-incubated mixture of (KaiB + KaiC-EE). Aliquots were taken from both tubes immediately after mixing (*t* = 6 h). Incubation continued at 30 ºC for an additional 6 h at the end of which aliquots were removed from each tube (*t* = 12 h). In the right panel, FAM-RpaA was mixed with SasA and KaiC-EE. An aliquot was removed immediately after mixing (*t* = 0 h). After incubating for 6 h at 30 ºC the sample was divided into two separate tubes. One tube received buffer and other tube received 3.5 µM KaiB after which aliquots were immediately taken (*t* = 6 h). Both samples were further incubated for 6 h at 30 ºC at the end of which aliquots were removed from each tube (*t* = 12 h). Time points for control reactions containing FAM-RpaA (Control-1), FAM-RpaA + SasA (Control-2) or FAM-RpaA + KaiC-EE + KaiB (Control-3) were also collected. **(B)** Aliquots were loaded onto 12.5% $Zn^{2+}$-Phos-tag$^{TM}$ SDS-PAGE for separation of RpaA phosphoforms. Coomassie blue-stained gel is in the top panel. Fluorescently labeled RpaA (FAM-RpaA) bands were visualized under UV transillumination (bottom panel). %RpaA~P was determined by densitometry.

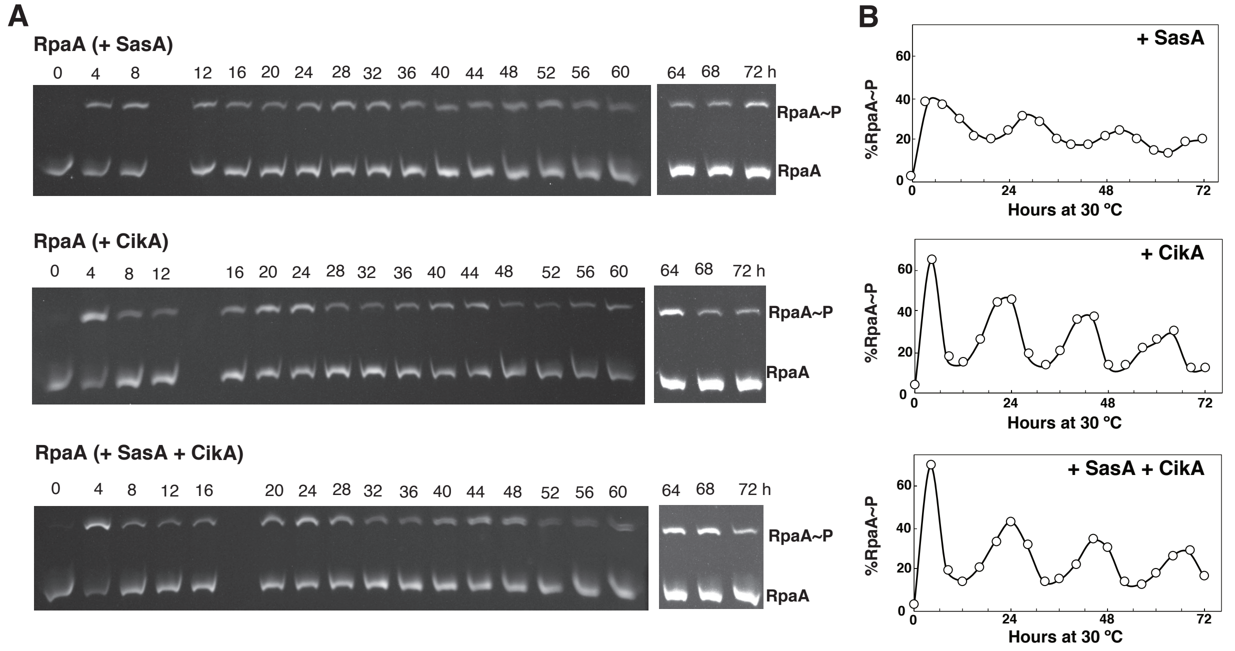

### Figure S5. Phos-tag$^{TM}$ SDS-PAGE for WT RpaA phosphorylation in vitro.

**(A)** Aliquots of reconstituted clock reactions containing KaiA, KaiB, KaiC, RpaA, DNA, and either SasA, CikA, or both were sampled every 4 h manually, subjected to 12.5% $Zn^{2+}$ Phos-tag$^{TM}$ SDS-PAGE, and visualized by UV-transillumination (see Materials and Methods for details). (**B**) Densitometry of the gels was performed by ImageJ (NIH) and band intensities were used to determine the level of RpaA phosphorylation at each time point using the equation

$$\%RpaA{\sim}P = 100\% * RpaA{\sim}P / (RpaA + RpaA{\sim}P)$$

Top, middle, and bottom panels are from IVC reactions that used SasA, CikA, and SasA + CikA.

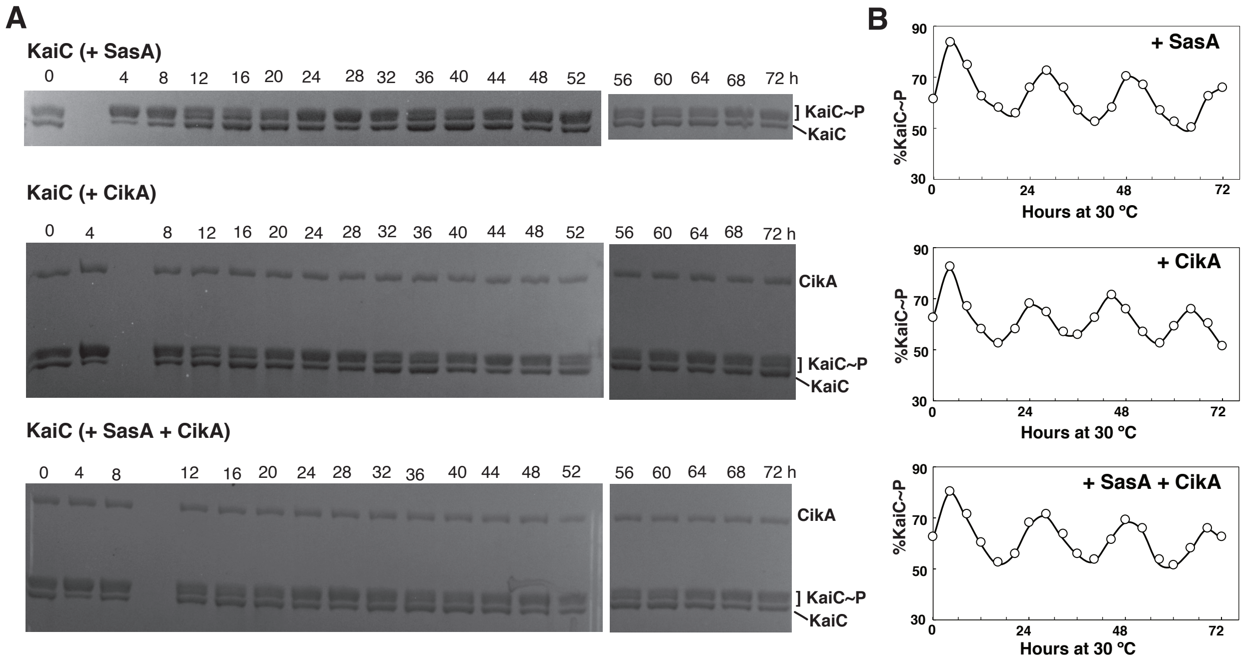

### Figure S6. SDS-PAGE of KaiC phosphorylation in vitro.

KaiC~P was measured in IVC reactions containing KaiA, KaiB, KaiC, RpaA, DNA, and either SasA (top), CikA (middle), or both SasA and CikA (bottom). (**A**) Aliquots were collected every 4 h, subjected to 7.5% SDS-PAGE, and stained with Instant Blue (Expedion) dye. The gels were analyzed by ImageJ software (NIH), where the sum of intensities of the top three bands were used to calculate the level of phosphorylated KaiC (KaiC~P). The bottom band is of unphosphorylated KaiC. (**B**) %KaiC~P = KaiC~P/total KaiC is plotted as function of time where top, middle, and bottom panels are from IVC reactions that used SasA, CikA, and SasA + CikA.

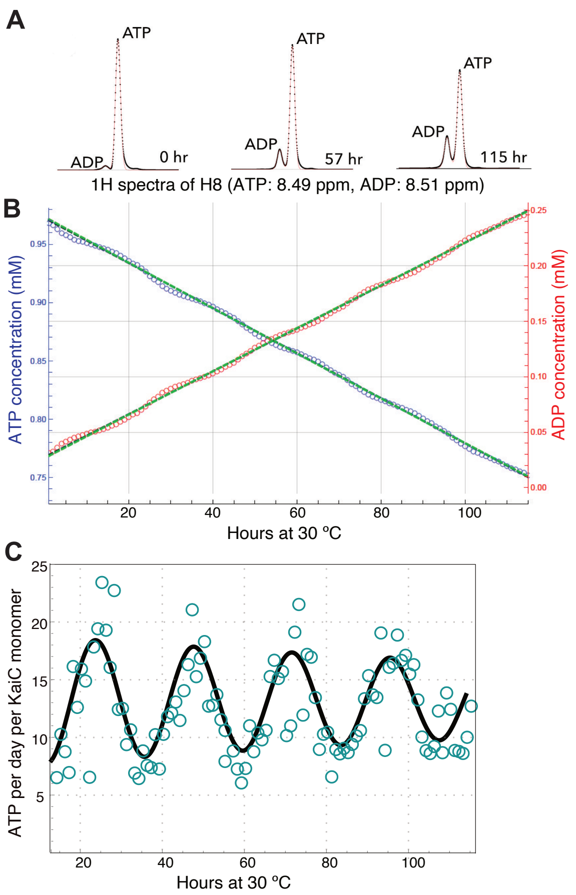

#### Figure S7. $^{1}$H NMR of ATP and ADP in a KaiABC oscillator reaction.

**(A)** H8 protons on the adenosine rings of ATP and ADP with chemical shifts of 8.49 ppm and 8.51 ppm, respectively, were monitored by $^{1}$H NMR as a function of reaction time. One-dimensional spectra were collected every hour and changes in peak intensities were measured. **(B)** Based on total nucleotide concentration of 1 mM, integrated peak intensities of ATP (blue) and ADP (red) at each time point were converted to concentrations in mM. The green lines are linear fits through the data. **(C)** ATPase activity is shown by teal circles and a cosine fit is shown in black. The total ATPase activity oscillated between 8 and 18 ATPs per day per KaiC monomer. During the first 12 hours, the sample was approaching a stable limit cycle and thus this data was not used in the fit. In a second measurement (data not shown) oscillations were between 7-13 ATPs per day per KaiC monomer.

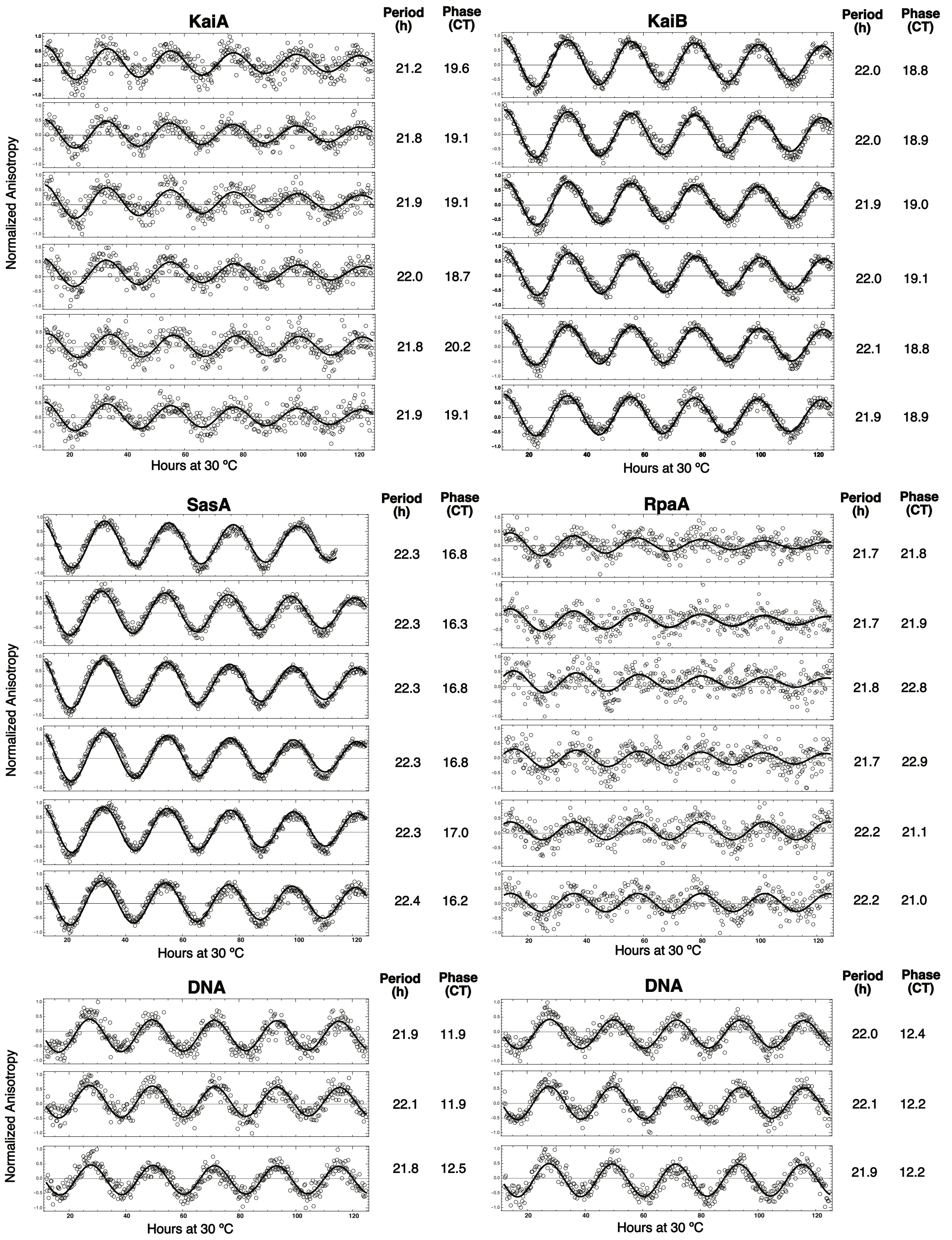

**Figure S8 (A). Fluorescence anisotropy data for IVC reactions containing SasA.** Normalized fluorescence anisotropy data are shown for three independent experiments, each containing two replicates for every fluorescence probe. Period and phase measurements were performed for each individual trace shown above, and average values of each repeat are reported in Fig.1B.

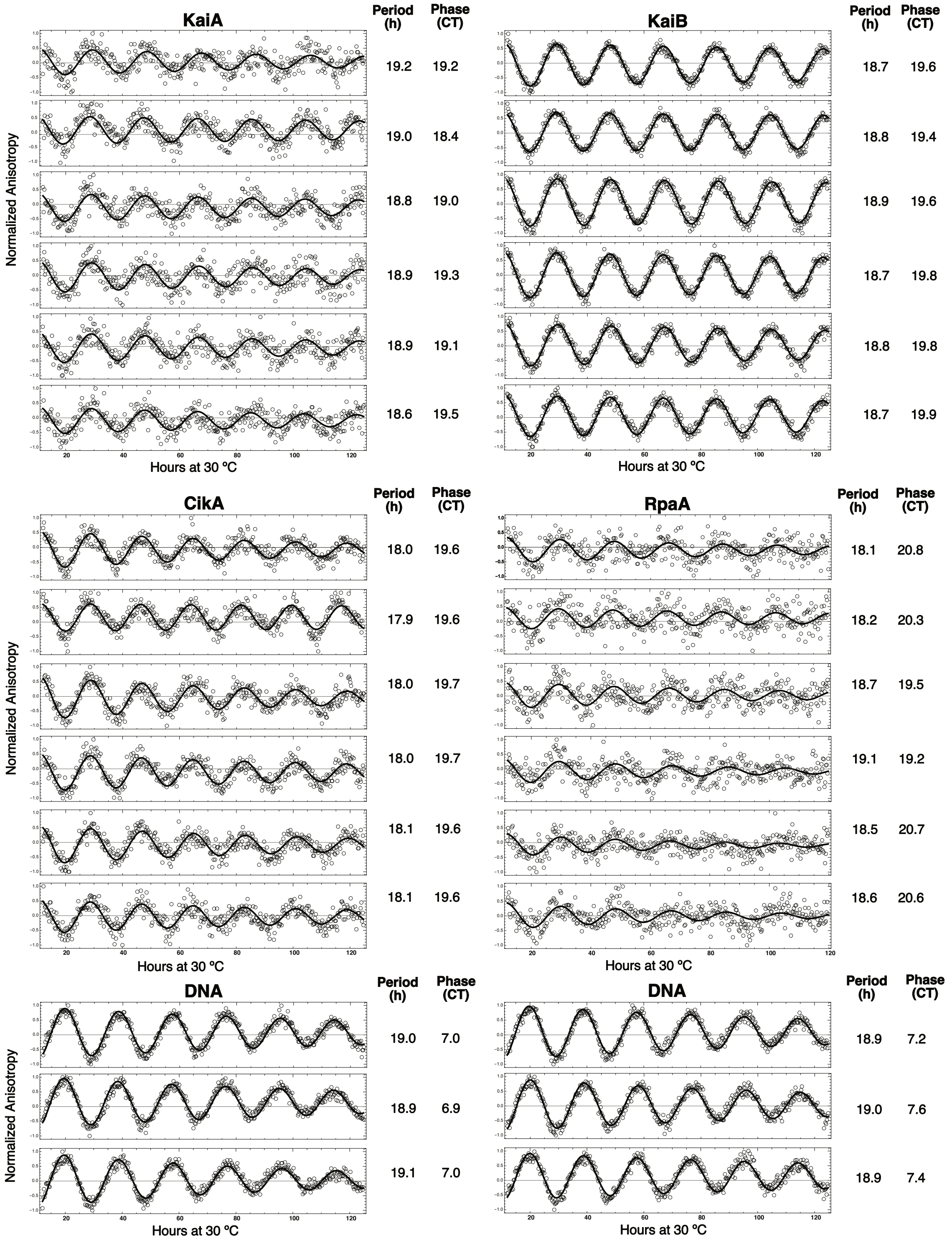

**Figure S8 (B). Fluorescence anisotropy data for IVC reactions containing CikA.** Normalized fluorescence anisotropy data are shown for three independent experiments, each containing two replicates for every fluorescence probe. Period and phase measurements were performed for each individual trace shown above, and average values of each repeat are reported in Fig. 2B.

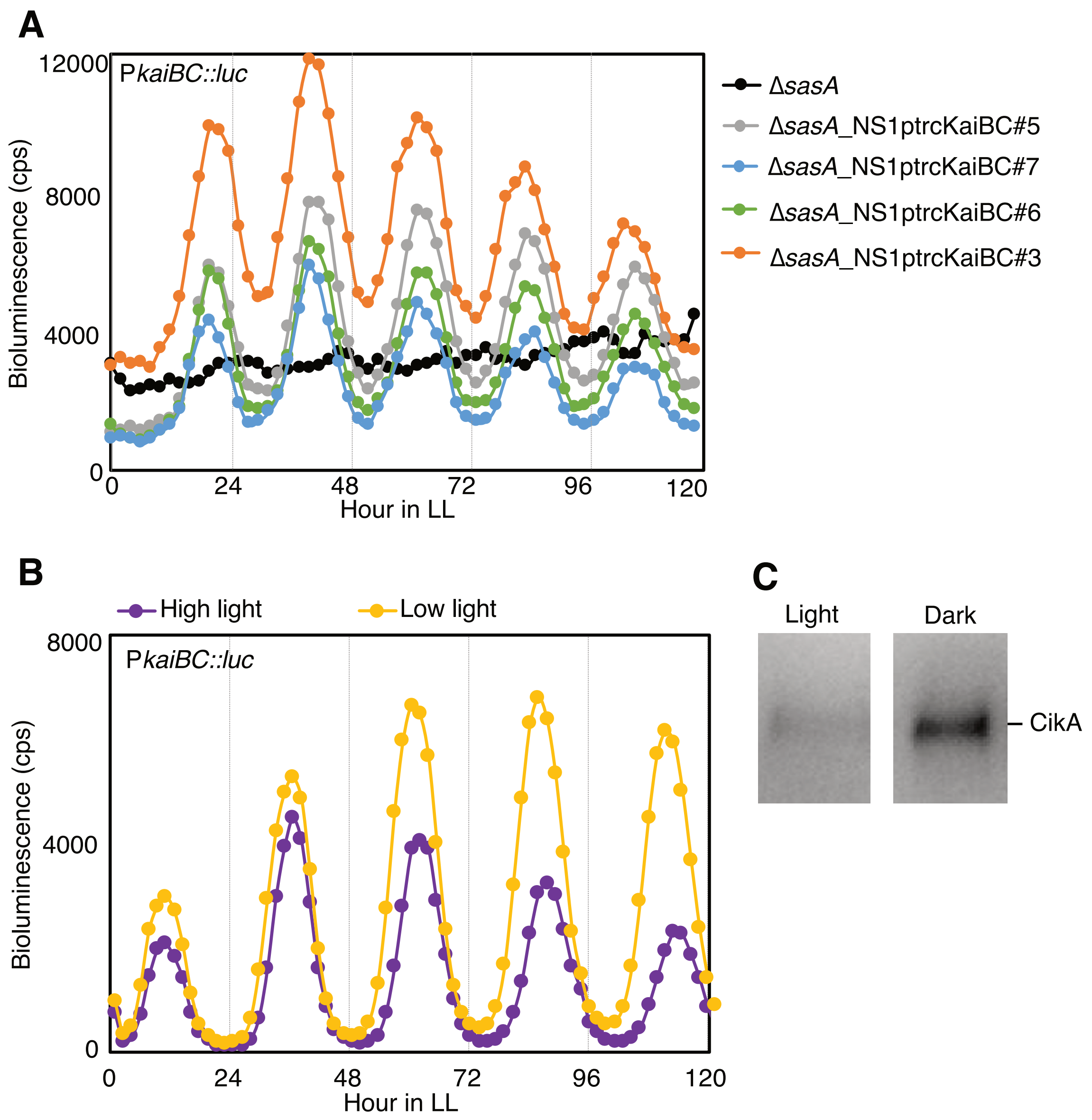

**Figure S9. Bioluminescence assays of different cyanobacterial strains.**

**(A)** Expression from P*kaiBC* monitored as bioluminescence from a P*kaiBC*::*luc* firefly luciferase reporter. Bioluminescence is nearly arrhythmic in a Δ*sasA* mutant (black). When *kaiBC* was driven by an *E. coli* promoter, P*trc*, in a Δ*sasA* background (*11*) to restore wild-type levels of KaiBC, rhythms were restored (traces from 4 replicates shown). **(B)** Rhythms of bioluminescence generated by P*kaiBC::luc* expression were monitored for entrained wild type (AMC541) in high light (purple) or low light (yellow) conditions. **(C)** CikA levels in AMC06 were probed using anti-CikA serum in cells grown in light or dark. Culture from one flask was split and kept under light or dark conditions for 6 h before harvesting.

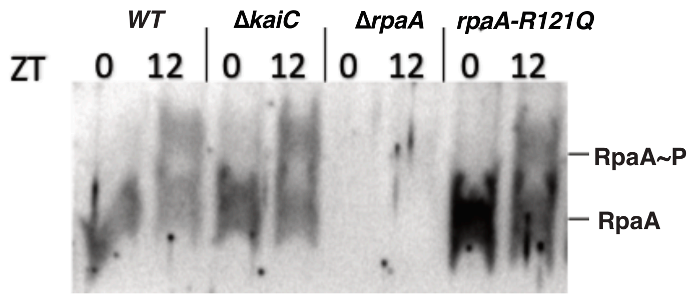

#### Figure S10. RpaA phosphorylation in vivo.

Phos-tag™ immunoblots reveal the level of RpaA phosphorylation in entrained wild-type, Δ*kaiC*, Δ*rpaA* and *rpaA-R121Q* cells at ZT = 0 (dawn) and ZT = 12 (dusk). Each lane contains 10 µg of total protein.

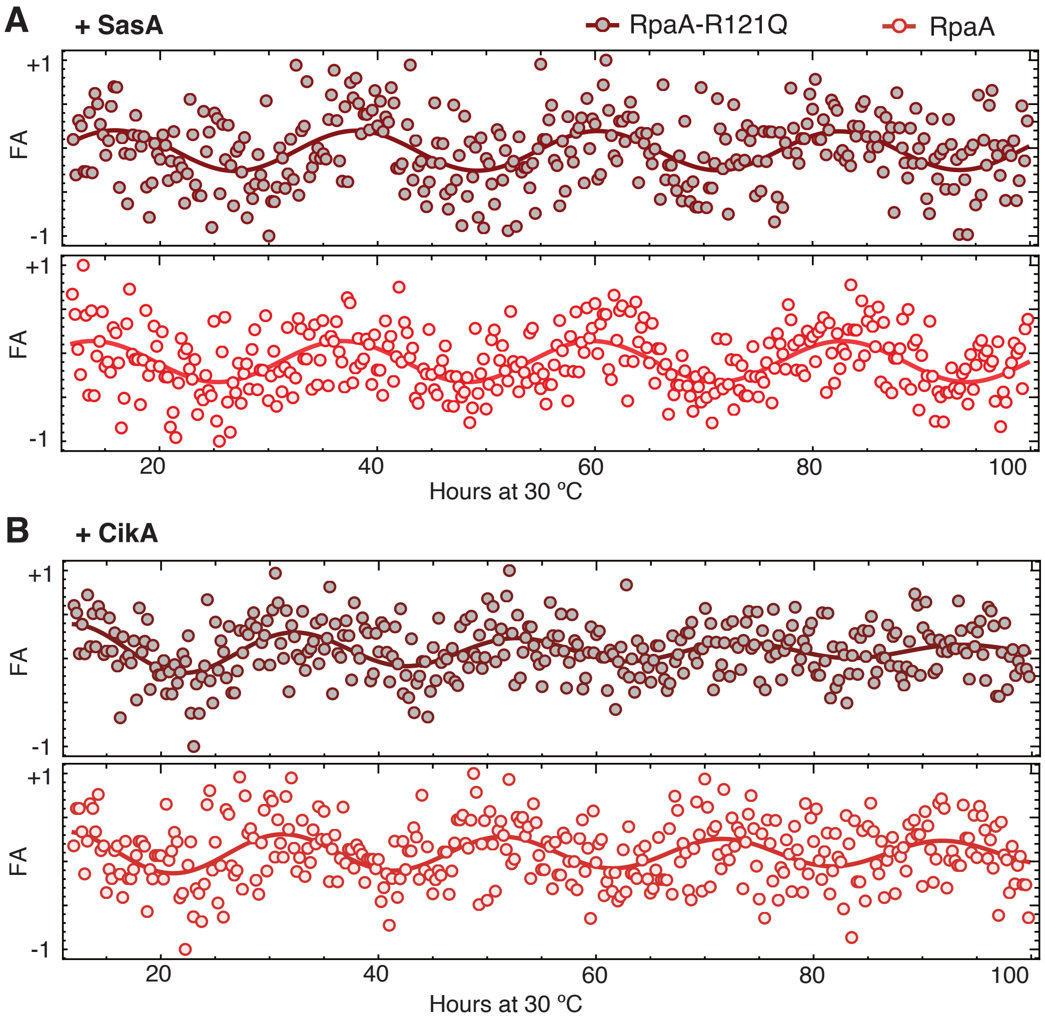

### Figure S11. RpaA and RpaA-R121Q fluorescence anisotropy in IVC.

Fluorescence anisotropy rhythms of labeled FAM-RpaA (red) and FAM-RpaA-R121Q (brown), measured in IVC containing KaiA, KaiB, KaiC and 0.65 µM of either (**A**) SasA or (**B**) CikA.

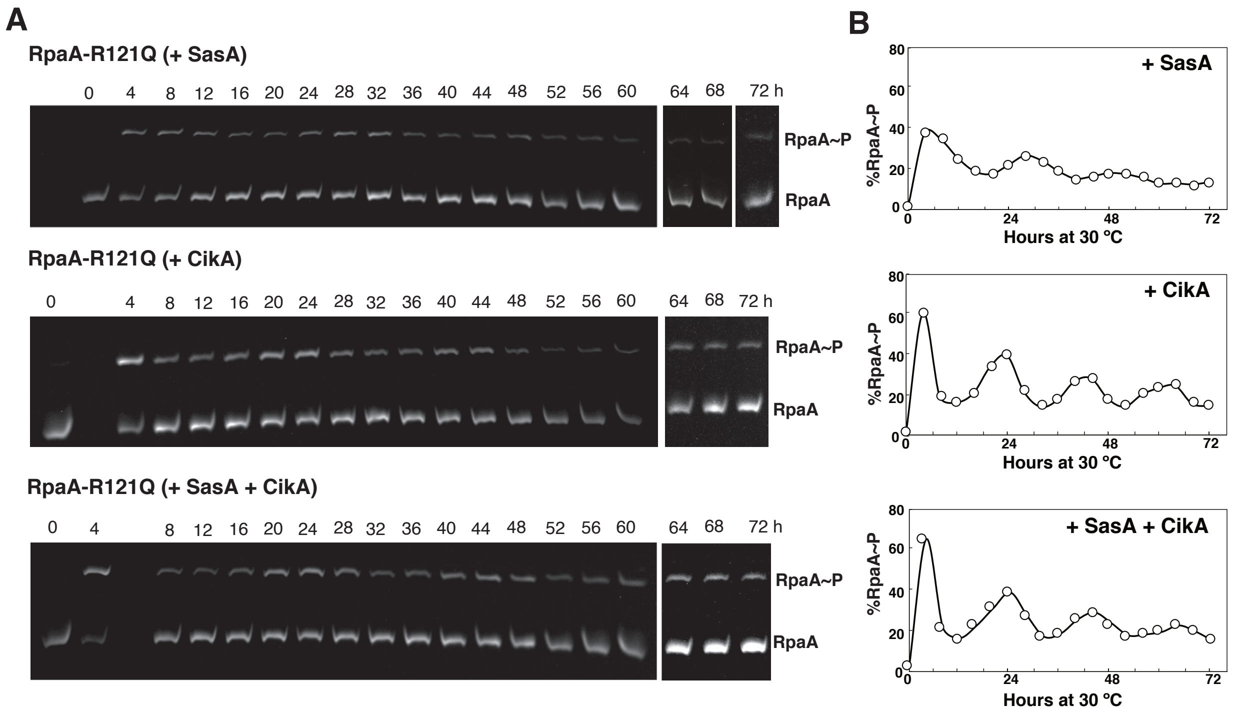

#### Figure S12. Phos-tag$^{TM}$ SDS-PAGE of RpaA-R121Q phosphorylation in vitro.

(**A**) Aliquots of reconstituted clock reactions containing KaiA, KaiB, KaiC, RpaA-R121Q, DNA, and either SasA (top), CikA (middle), or both (bottom) were sampled every 4 h manually, subjected to 12.5% $Zn^{2+}$ Phos-tag$^{TM}$ SDS-PAGE, and visualized by UV-transillumination (see Materials and Methods for details). (**B**) Densitometry of the gels was performed by ImageJ (NIH) and %RpaA~P = 100% * RpaA~P / (RpaA + RpaA~P) is plotted as function of time.
